# PTK2 regulates the UPS impairment via p62 phosphorylation in TDP-43 proteinopathy

**DOI:** 10.1101/355446

**Authors:** Shinrye Lee, Yu-Mi Jeon, Seyeon Kim, Younghwi Kwon, Myungjin Jo, You-Na Jang, Seongsoo Lee, Jaekwang Kim, Sang Ryong Kim, Kea Joo Lee, Sung Bae Lee, Kiyoung Kim, Hyung-Jun Kim

## Abstract

TDP-43 proteinopathy is a common feature in a variety of neurodegenerative disorders including Amyotrophic lateral sclerosis (ALS) cases, Frontotemporal lobar degeneration (FTLD), and Alzheimer’s disease. However, the molecular mechanisms underlying TDP-43-induced neurotoxicity are largely unknown. In this study, we demonstrated that TDP-43 proteinopathy induces impairment in ubiquitin-proteasome system (UPS) evidenced by an accumulation of ubiquitinated proteins and reduction of proteasome activity in neuronal cells. Through kinase inhibitor screening, we identified PTK2 as a suppressor of neurotoxicity induced by UPS impairment. Importantly, PTK2 inhibition significantly reduces ubiquitin aggregates and attenuated TDP-43-induced cytotoxicity in *Drosophila* model of TDP-43 proteinopathy. We further identified that phosphorylation of p62 at serine 403 (p-p62^S403^), a key component in the autophagic degradation of poly-ubiquitinated proteins, is increased upon TDP-43 overexpression and dependent on activation of PTK2 in neuronal cells. Moreover, expressing a non-phosphorylated form of p62 (p62^S403A^) significantly represses accumulation of polyubiquitinated proteins and neurotoxicity induced by TDP-43 overexpression in neuronal cells. In addition, inhibition of TBK1, a kinase which phosphorylates S403 of p62, ameliorates neurotoxicity upon UPS impairment in neuronal cells. Taken together, our data suggest that activation of PTK2-TBK1-p62 axis plays a critical role in the pathogenesis of TDP-43 by regulating neurotoxicity induced by UPS impairment. Therefore, targeting PTK2-TBK1-p62 axis may represent a novel therapeutic intervention for neurodegenerative diseases with TDP-43 proteinopathy.

## Introduction

Amyotrophic lateral sclerosis (ALS) is the most common type of motor neuron disease characterized by the progressive degeneration of motor neurons in the brain and the spinal cord (Al-Chalabi et al, 2012; Cleveland & Rothstein, 2001). ALS is a fatal neurodegenerative disease with no cure. Currently, there is only one FDA-approved drug for ALS. However, it is known to extend lifespan by only a few months (Costa et al, 2010). Therefore, there is an urgent demand for the development of effective treatments for ALS.

Intracellular accumulation of TDP-43 is a pathological hallmark of ALS in the majority of cases (James et al, 2016; Josephs et al, 2017; Liscic et al, 2008; Scotter et al, 2015). TDP-43 is a highly conserved nuclear RNA binding protein. So far, over fifty missense mutations of TDP-43 are linked to sporadic and familial ALS (Buratti, 2015; Scotter et al, 2015). TDP-43-induced neurotoxicity is currently well recognized to contribute to the pathology of ALS and other neurodegenerative diseases linked to the deposition of TDP-43 called TDP-43 proteinopathy, such as Frontotemporal lobar degeneration (FTLD) and Alzheimer’s disease (James et al, 2016; Meriggioli & Kordower, 2016; Scotter et al, 2015). However, the neuropatholical mechanisms of TDP-43 proteinopathy is largely unknown. To get further insights into the pathogeneses of TDP-43 proteinopathies, it is critical to understand how TDP-43 deposition leads to the neurotoxicity.

Of note, the most prominent pathological hallmark of TDP-43 proteinopathies is the presence of intracellular ubiquitin-positive inclusions in neurons (Bendotti et al, 2012; Neumann et al, 2006). Ubiquitinated proteins are known to be rapidly degraded by the proteasome in healthy neurons (Ross et al, 2015). These findings raise the possibility that the accumulation of ubiquitin-positive proteinaceous inclusions in degenerating neurons has been attributed to impairment of the ubiquitin proteasome system (UPS) in TDP-43 proteinopathy. Interestingly, several genes associated with ALS, such as *SQSTM1*, *VCP*, *OPTN*, *TBK1* and *UBQLN2*, are directly linked to the degradation of ubiquitinated proteins, suggesting that impairment of UPS may be a core pathological mechanism of ALS (Lee et al, 2015; Renton et al, 2014). These all together raise the possibility that UPS may play a crucial role in TDP-43 proteinopathy. However, it has never been answered whether UPS impairment is involved in neurotoxicity of TDP-43 accumulation.

Notably, *Drosophila* models of TDP-43 proteinopathy recapitulate many key pathological features of ALS, such as deposition of ubiquitin-positive aggregates, the progressive motility deficits, and shortened lifespan (Casci & Pandey, 2015; Joyce et al, 2011). In this study, we investigated the potential role of UPS in TDP-43-induced neuropathology by utilizing *Drosophila* model as well as mammalian cellular models of TDP-43 proteinopathy. We demonstrated that proteasome activity is significantly reduced by TDP-43 accumulation. Moreover, we identified that PTK2 plays a critical role in deposition of ubiquitinated aggregates and neurotoxicity induced by UPS impairment through regulating TBK1-p62 pathway in TDP-43 proteinopathy.

## Methods

### Reagents and Antibodies

The following reagents were purchased as indicated: dimethyl sulfoxide (DMSO) and mifepristone (RU-486) [Sigma]; MG132 [Calbiochem/Merck-Millipore]; PTK2 inhibitor (PF573228) and CK2 inhibitor (DMAT) [APExBIO]; ULK1 inhibitor (MRT68921) [Selleckchem]; TBK1 inhibitor (BX 795) [Axon Medchem]. The following antibodies were used for immunoblotting: rabbit anti-phospho-PTK2 (Tyr397) (catalog no. 3283), rabbit anti-PTK2 (catalog no. 3285), and HRP-conjugated anti-alpha-tubulin (catalog no. 9099) [Cell Signaling Technology]; mouse anti-TBK1 (catalog no. ab40676); HRP-conjugated anti-mouse IgM (catalog no. ab97230), and rabbit anti-beta-actin (catalog no. ab16039) [Abcam]; mouse anti-Poly-ubiquitin (FK1) (catalog no. BML-PW8805), [Enzo Life Science]; rabbit anti-TDP-43 (catalog no. 10782-2-AP) [Proteintech]; mouse anti-GFP (catalog no. 632380) [Clontech]; rabbit anti-p62/SQSTM1 (catalog no. P0067) [Sigma]; mouse anti-poly-ubiquitin (FK2) (catalog no. 04-263) [Calbiochem/Merck-Millipore]; rabbit anti-phospho-p62 (Ser403) (catalog no. GTX128171) [GeneTex]; mouse anti-turboGFP (catalog no. TA150041) [Origene]; HRP-conjugated anti-rabbit IgG (catalog no. sc-2004), and HRP-conjugated anti-mouse IgG (catalog no. sc-2005) [Santa Cruz]. The following antibodies were used for immunocytochemistry: rabbit anti-cleaved caspase-3 (catalog no. 9611) [Cell Signaling Technology]; mouse anti-poly-ubiquitin (FK1) (catalog no. BML-PW8805) [Enzo Life Science]; rabbit anti-phospho-PTK2 (Tyr397) (catalog no. ab81298) [Abcam]; mouse anti-DDK (catalog no. TA50011) [Origene]; and rabbit anti-phospho-p62 (Ser403) (catalog no. GTX128171) [GeneTex]. The following antibodies were used for immunohistochemistry: rat anti-elav (catalog no. RAT-ELAV-7) [DSHB] and mouse anti-poly-ubiquitin (FK1) (catalog no. BML-PW8805) [Enzo Life Science].

### Cell lines

The SH-SY5Y human neuroblastoma cell line and Neuro-2a (N2a) mouse neuroblastoma cell line were maintained in DMEM (Life Technologies) supplemented with 10% heat-inactivated fetal bovine serum (FBS) (Gibco) and penicillin-streptomycin (50 μg/ml) (Gibco). Primary cultures of cerebral cortical neurons were prepared from 16-day embryonic mice as described previously (Araki et al, 2000; Enokido et al, 1992). Briefly, mouse embryos were decapitated, and the brains were rapidly removed and placed in a culture dish containing HBSS (Gibco). Cortices were isolated, transferred to a conical tube and washed twice in HBSS (Gibco). Cortical tissues were enzymatically digested by papain (20 units/ml) (Worthington Biochemical Corporation) and DNase I (0.005 %) for 30 min at 37 °C. The tissues were mechanically dissociated by gently pipetting up and down. The cortical cells were centrifuged at 800 rpm for 10 min at room temperature. Then, the dissociated cells were seeded onto plates coated with poly-D-lysine (Sigma-Aldrich) in neurobasal media containing 2 mM glutamine (Gibco), N2 supplement (Gibco), B27 supplement (Gibco), and penicillin-streptomycin (Gibco). Culture media were changed initially after 5 days and then changed every 3 days. All experiments with primary neurons were performed at DIV 14-21 days.

### Transfection

N2a cells were plated in six-well plate and transfected with 4 μg of pCMV6-AC-TDP-43-GFP expressing green fluorescent protein (GFP)-tagged human TDP-43 per well by using Lipofectamine 3000 reagent (Invitrogen) according to the manufacturer’s guide. An empty pCMV6-AC-GFP plasmid was used as a negative control. For siRNA transfection, cells were transfected with control siRNA (Santa Cruz; sc-37007), mouse *p62* siRNA (Santa Cruz; sc-29828), mouse *ptk2* siRNA (Santa Cruz; sc-35353), or mouse *tbk1* siRNA (Santa Cruz; sc-39059) by using Lipofectamine 3000 reagent. 48 h post-transfection, knockdown of target proteins was confirmed by immunoblot analysis.

### Immunostaining

For immunocytochemistry, cells were fixed in 4 % paraformaldehyde in PBS for 30 min. The following antibodies were used for immunostaining as indicated: mouse anti-poly-ubiquitin (FK1), rabbit anti-LC3, rabbit anti-PTK2 (Tyr397), rabbit anti-cleaved caspase-3, mouse anti-DDK, rabbit anti-phospho-p62 (Ser403), and rabbit anti-p62. Alexa-594 conjugated goat anti-rabbit IgG, Alexa-488 conjugated goat anti-mouse IgG, and Alexa-594 conjugated goat anti-mouse IgM (Jackson labs) were used as secondary antibodies as indicated. Antibodies were incubated in PBS with 0.3 % Triton and 1 % BSA. Then, samples were mounted and observed with a fluorescence microscope. For immunohistochemistry, adult flies were dissected in PBS and fixed in 4 % paraformaldehyde in PBS for 30 min. After incubating together with rat anti-elav (7E8A10) and mouse anti-poly-ubiquitin (FK1) in PBS with 0.3 % Triton and 5 % normal goat serum, the sections were incubated with Alexa-488 conjugated goat anti-rat IgG and Alexa-594 conjugated goat anti-mouse IgM (Jackson labs). All antibodies were diluted in PBS with 0.3 % Triton and 5 % normal goat serum. The samples were mounted and observed with fluorescence microscope or confocal microscope

### Proteasome activity assay

N2a cells were transfected with pCMV6-AC-GFP or pCMV6-AC-TDP-43-GFP for 3 days and then the cells expressing GFP were sorted using FACS. Cells harvested were incubated in Lysis Buffer (25 mM HEPES (pH 7.4), 10 % glycerol, 5 mM MgCl_2_, 1 mM ATP, 1 mM DTT). The lysates were centrifuged at 13,000 × g for 30 min at 4°C. 100 μg of total protein were used per proteasome purification by using 26S proteasome purification kit according to the manufacturer’s guide (Enzo Life Science). The lysates were incubated with matrices (Enzo Life Science) for overnight at 4 °C. 20S proteasome activity were measured using the 20S proteasome activity Kit (Enzo Life Science) according to the manufacturer’s guide. Briefly, 20 μg of purified lysates were incubated with inhibitor and enzyme matrix (Enzo Life Science) for 10 min at RT. After adding 10 μL of fluorescently labeled substrate (trypsin-like; Boc-LSTR-AMC, Bachem and chymotrypsin-like; Suc-LLVY-AMC; caspase-like; Z-nLPnLD-AMC) to each well, the fluorescence signals were measured at 10-min intervals for 1 h using a FlexStation 3 microplate reader (set to 37 °C, Molecular Devices) at an excitation wavelength of 360 nm and an emission wavelength of 460 nm.

### Preparation of soluble and insoluble cell extracts

Cells were homogenized in RIPA buffer with protease and phosphatase inhibitor cocktails. Soluble and insoluble fractions in 1 % Triton X-100 were obtained by centrifugation at 100,000 × g for 30 min at 4 °C. Supernatants containing the soluble factions were harvested and the pellets for insoluble fractions were solubilized in a 2 % SDS detergent RIPA buffer. After sonication, the cell lysates were mixed with NuPAGE LDS Sample buffer containing reducing agent (Invitrogen) and then boiled at 95 °C for 5 min.

### Immunoblot analysis

For total protein extraction, either cells or 20 adult fly heads were homogenized in RIPA buffer (Cell signaling Technology) or 1 × LDS sample buffer (Invitrogen) with protease and phosphatase inhibitor cocktails (Roche). The protein concentration of the cell lysates was determined by BCA protein assay (Pierce). Next, the protein extracts were mixed with NuPAGE LDS Sample buffer containing reducing agent and then boiled at 95 °C for 5 min. An equal amount of proteins from each sample were separated on NuPAGE 4-12 % Bis-Tris gels (Novex) or NuPAGE 3-8 % Tris-Acetate gels (Novex) and transferred to a polyvinylidene difluoride (PVDF; Novex) membrane. After blocking membranes with 5 % skim milk in TBS with 0.025 % Tween 20, blots were probed with antibodies as indicated and detected with an ECL prime kit (Amersham Biosciences). Samples from three independent experiments were used and the relative expression levels were determined using a Fusion-FX (Viber Lourmat).

### Cytotoxicity Assay

SH-SY5Y cells (8 × 10^4^ cells/ml), N2a cells (5 × 10^4^ cells/ml), or primary cortical neuron cells (10 × 10^4^ cells/ml) were grown in 96-well plates and treated with MG132 or inhibitors as indicated for 24 h. DMSO was used as a negative control. To measure cytotoxicity, the Cell Counting Kit-8 (CCK-8; Enzo Life Science) was used according to the manufacturer’s guide. Briefly, 10 μL of CCK-8 reagent was added into each well and the plate was incubated at 37 °C for 2 h. The absorbance at 450 nm was measured by using a microplate reader (Tecan). Cell viability was expressed as a percentage of control. All experiments were performed in triplicate.

### Cyto-ID autophagy analysis

To detect autophagic vesicles, the Cyto-ID^®^ Green Autophagy Kit (Enzo Life Science) was used according to the manufacturer’s guide. Briefly, 1 μL of Cyto-ID dye was added into each well of a 24-well plate. Then, the plate was incubated at 37 °C for 30 min. Cells were fixed in 4 % paraformaldehyde in PBS for 30 min. All reagents were diluted in 1 × Assay buffer with 2 % FBS. The samples were mounted and observed with a microscope.

### Stable transfection

N2a cells in six-well plates were transfected with 4 μg of p62 or p62-S403A cDNA by using Lipofectamine 3000 reagent (Invitrogen). An empty pCMV6-Myc-DDK vector was used as a negative control. Stable transfectants were selected in the presence of G418 (800 μg/ml). The expression of transgenes was confirmed by immunoblot and immunocytochemistry analysis.

### Quantitative RT-PCR

RNA was extracted from cells by using TRIzol reagent (Life Technologies). RNA cleanup was performed using the RNeasy Mini Kit (Qiagen) according to the manufacturer’s instructions. cDNA synthesis was performed at 37 °C for 120 min with 100 ng of RNA using a High Capacity cDNA Reverse Transcription kit (Applied Biosystems). Quantitative RT-PCR was performed using the one-step SYBR^®^ PrimeScript™ RT-PCR kit (Perfect Real Time; Takara Bio Inc.) according to the manufacturer’s instructions, followed by detection using an Applied Biosystems 7500 Real-Time PCR system (Applied Biosystems). GAPDH was used as an internal control. The 2^−ΔΔCt^ method was used to calculate relative changes in gene expression, as determined by real time PCR experiments (Livak & Schmittgen, 2001).

### Fly strains

Drosophila stocks were maintained on standard cornmeal agar media at 24 °C unless otherwise noted. UAS-TDP-43 and UAS-ATXN2-32Q were described previously (Kim et al, 2014). UAS-CL1-GFP has been described previously (Pandey et al, 2007). All other stocks were from The Bloomington stock center.

### Climbing assays

Adult males (0 to 1 day old) were separated and transferred into experimental vials containing fly media or paper mixed with or without RU-486 (in ethanol, 40 μg/ml) at a density of 25 flies per vial (n > 100). The number of dead flies was scored daily, and flies were transferred to fresh media or paper every other day. Adult locomotor function was assessed by a previously described method (Feany & Bender, 2000) with 125 flies per genotype per time point in all experiments. Experiments were repeated twice to assure consistent results.

### Statistical analyses

Data were analyzed by Student’s *t*-test (Vassar Stats, www.vassarstats.net), one-way ANOVA, or two-way ANOVA test depending on comparison variables with post-hoc analysis as indicated (GraphPad Prism Software, La Jolla, CA). Differences were considered significant when p<0.05 and are indicated as follows: *p<0.05; **p*0.005; ***p<0.001; n.s., not significant.

## Results

### TDP-43 overexpression induces UPS impairment

To see if TDP-43 overexpression impairs UPS in neuronal cells, we first transfected GFP-tagged human TDP-43 or negative control to mouse neuronal N2a cells (Fig. 1A). Consistent with previous findings (Bendotti et al, 2012), TDP-43 overexpression dramatically increased accumulation of poly-ubiquitinated proteins in N2a cells compared to control (Fig. 1B). TDP-43 overexpression markedly increased the levels of poly-ubiquitinated proteins in insoluble fractions, whereas it mildly increased the levels of poly-ubiquitinated proteins in soluble fractions (Fig. 1C). To examine whether TDP-43 overexpression affects proteasome activity, we measured activities of 26S proteasomes purified from cell lysates. Interestingly, trypsin-like activity of the proteasomes from TDP-43-overexpressing cells was significantly decreased than that from control cells, whereas chymotrypsin-like activity was not altered by TDP-43 overexpression (Fig. 1D). It is well known that autophagy-lysosomal pathway (ALP) is activated upon UPS impairment as a compensatory mechanism (Wang & Wang, 2015). In line with this, we also observed that TDP-43 overexpression activated ALP evidenced by increased levels of LC3 puncta and LC3-II/LC3-I ratio in N2a cells (Fig. S1A and B) compared to control, further supporting that TDP-43 overexpression impairs UPS in neuronal cells. Taken together, our data demonstrated that TDP-43 overexpression decreases proteasome activity and thereby increases poly-ubiquitinated proteins in neuronal cells.

**Figure 1.**
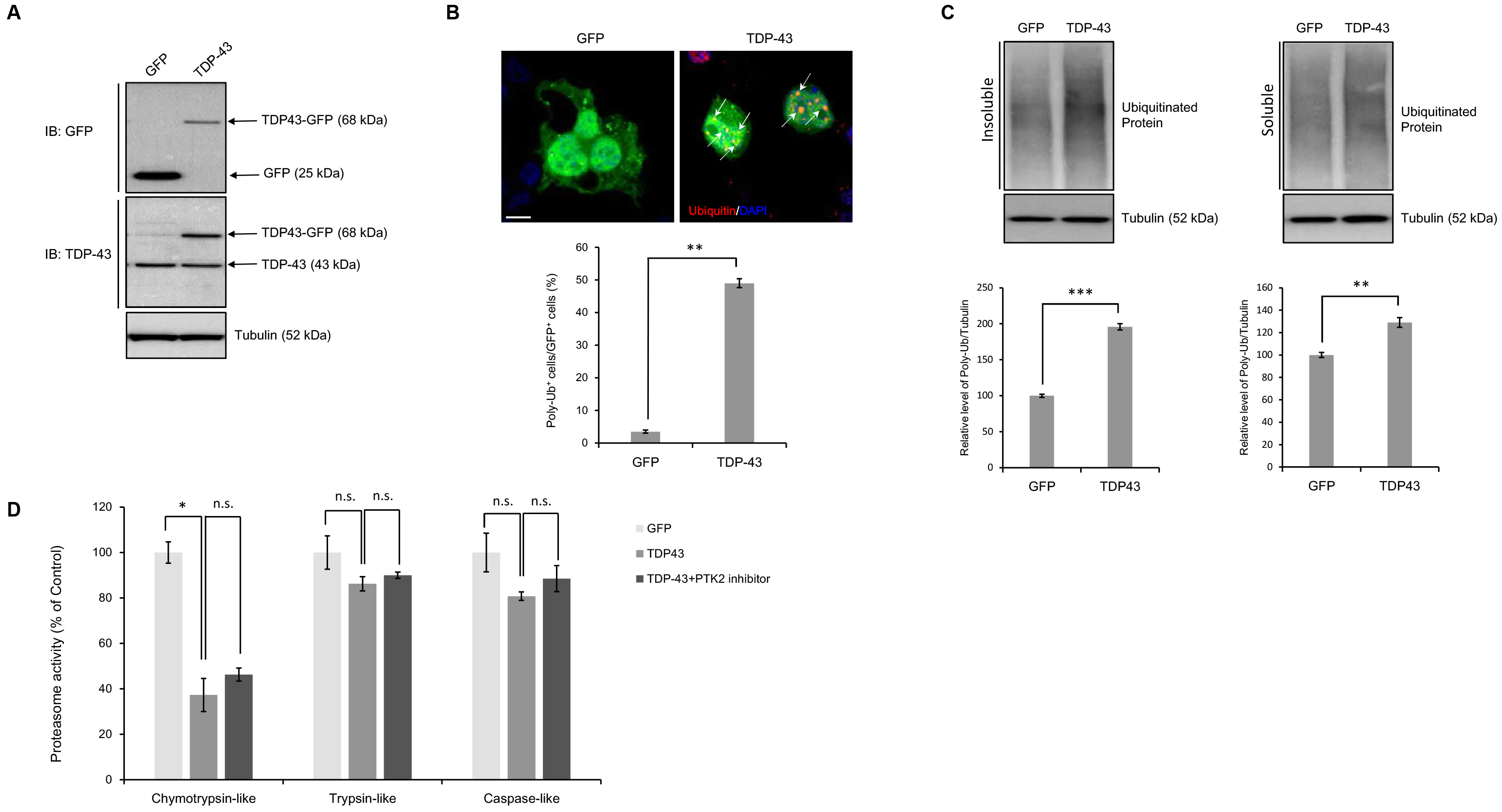
Overexpression of TDP-43 induced UPS impairment in neuronal cells. **A-C**, N2a cells were transiently transfected with the control vector (pCMV6-AC-GFP) or TDP-43 expression construct (pCMV6-AC-TDP-43-GFP) for 3 days. Immunoblotting or immunocytochemistry was performed thereafter. **(A)** Immunoblotting for GFP and TDP-43 protein expression. Quantification of the immunoblots from 3-5 independent experiments and normalized to tubulin. **(B)** Immunocytochemistry was subsequently performed to detect ubiquitin (red) or DAPI (nuclei; blue). Quantification of the percentage of poly-ubiquitin-positive staining in GFP-positive cells (*lower*). Arrowheads indicate the co-localization of the poly-ubiquitin with TDP-43-GFP-positive cells. Scale bars, 20 μm. Data are presented as the means ± SD of 3 independent experiments. **p<0.005 (Student’s *t*-test). Scale bars, 10 μm. **(C)** Cells were fractionated into the supernatant (soluble) and pellet (insoluble). Western blot analysis of poly-ubiquitinated proteins from the soluble and insoluble fractions. Quantification of immunoblots from 3-5 independent experiments and normalized to tubulin. Data are presented as the means ± SD. **p<0.005, ***p<0.001 (Student’s *t*-test). **(D)** N2a cells were transiently transfected with the control vector (pCMV6-AC-GFP) or TDP-43 expression construct (pCMV6-AC-TDP-43-GFP) for 2 days, subsequently treated with PTK2 inhibitor (5 μm) for 24 h. Chymotrypsin-like, Trypsin-like, or Caspase-like proteasome activity was subsequently performed to measure of fluorometric substrate. Slopes relative to that of control are shown. Data are presented as the means ± SD (n=3). *p<0.05, n.s. not significant (one-way ANOVA with Bonferroni multiple comparison test).

### PTK2 inhibition ameliorates neurotoxicity induced by TDP-43 overexpression

Given that accumulation of poly-ubiquitinated proteins exerts a cytotoxic effects on cells, we hypothesized that UPS impairment may mediate neurotoxicity in TDP-43 proteinopathy, which may be attenuated by modulating deficits in UPS. To address this hypothesis, we screened molecules which regulate cytotoxicity induced by blocking UPS with MG132 through kinase inhibitor screening. We identified PF573228 as a potent suppressor of MG132-induced toxicity (Fig. S2). PF573228 is a PTK2-specific inhibitor (Slack-Davis et al, 2007; Wilson et al, 2014). The levels of activated PTK2 phosphorylated at Tyr397 (p-PTK2^Y397^) was significantly increased upon MG132 treatment in human neuronal SH-SY5Y cells compared to control (Fig. S2A). Inhibiting PTK2 with PF573228 significantly decreased the levels of p-PTK2 ^Y397^ (Fig. S2A) and poly-ubiquitinated proteins (Fig. S2B and C) and cytotoxicity induced by MG132 in SH-SY5Y cells (Fig. S2D). Similar results were observed in mouse neuronal N2a cells and primary cortical neurons (Fig. S2E-H). Inhibiting PTK2 by PTK2-specific siRNA similarly attenuated MG132-induced cytotoxicity, ruling out the possibility of off-target effects of PF573228 (Fig. S3). To confirm these results, we used lactacystin as a specific UPS inhibitor (Yang et al, 2008). Consistently, lactacystin treatment also increased phpspho-PTK2 and PTK2 inhibition significantly suppressed lactacystin induced toxicity in N2a and SH-SY5Y cells (Fig. S4).

Given that PTK2 regulates cytotoxicity induced by UPS impairment, we next investigated whether inhibiting PTK2 ameliorates neurotoxicity induced by overexpressing TDP-43. We first assessed if TDP-43 overexpression increases the levels of p-PTK2^Y397^. TDP-43 overexpression markedly increased p-PTK2 ^Y397^ levels in N2a cells compared to control (Fig. 2A). Inhibiting PTK2 by either siRNA or PF573228 dramatically decreased accumulation of poly-ubiquitinated proteins by TDP-43 overexpression in N2a cells (Fig. 2B). To see if PTK2 regulates neurotoxicity induced by TDP-43 overexpression, we monitored the levels of caspase-3 after inhibiting PTK2 in N2a cells expressing TDP-43. Importantly, inhibiting PTK2 dramatically decreased an induction of caspase-3 by TDP-43 overexpression in N2a cells compared to controls (Fig. 2C), indicating that PTK2 inhibition strongly suppresses neurotoxicity in cellular model of TDP-43 proteinopathy.

**Figure 2.**
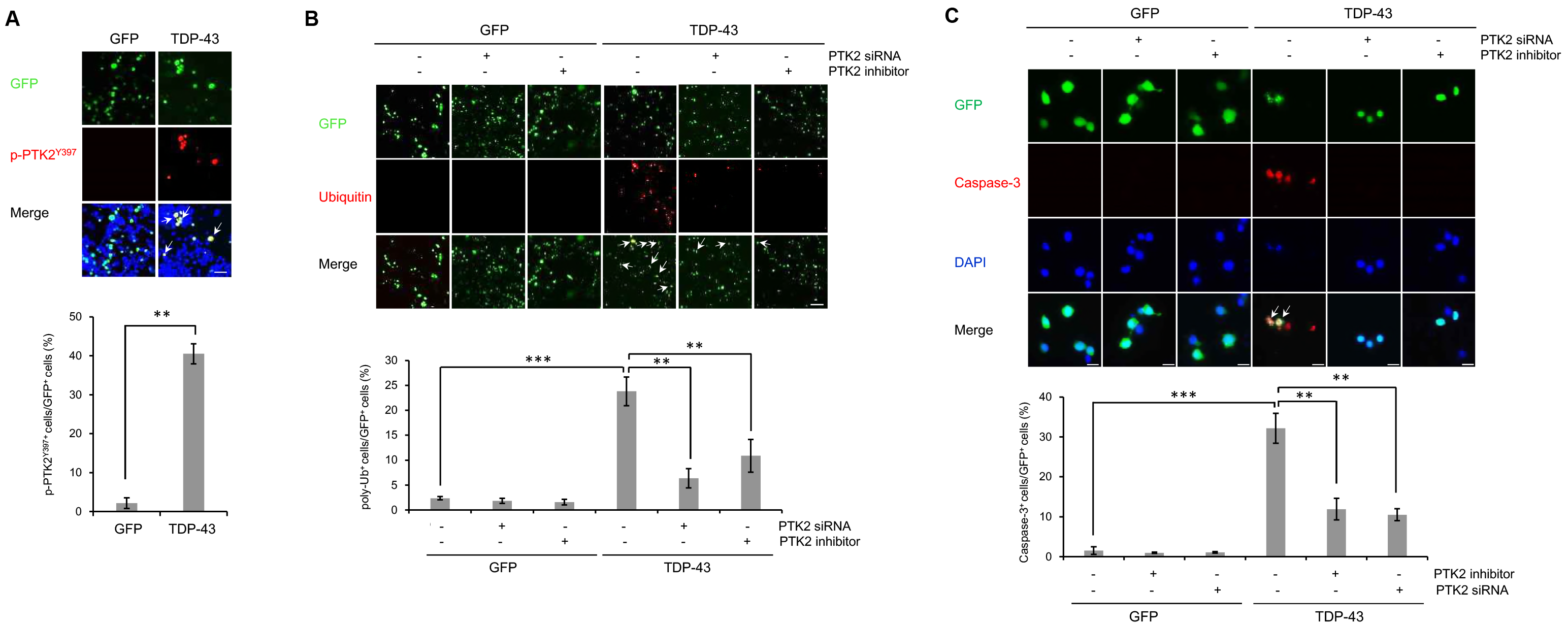
Inhibition of PTK2 restores TDP-43-induced UPS impairment and neuronal toxicity. **(A-C)** N2a cells were transiently transfected with the control vector (pCMV6-AC-GFP) or TDP-43 expression construct (pCMV6-AC-TDP-43-GFP) for 3 days. Cells overexpressing GFP or TDP-43-GFP were treated with *ptk2*-specific siRNA (20 nM) or a PTK2 inhibitor (5 μm) for 24 h. Immunocytochemistry was performed thereafter. (A) Immunocytochemistry was subsequently performed to detect PTK2 phosphorylation (p-PTK2^Y397^; red). Quantification of the percentage of p-PTK2^Y397^-positive cells in GFP-positive cells (*lower*). Arrowheads indicate the co-localization of the p-PTK2^Y397^ with TDP-43-GFP-positive cells. Data are presented as the means ± SD of 3 independent experiments. **p<0.005 (Student’s *t*-test). Scale bars, 200 μm. **(B)** Immunocytochemistry was subsequently performed to detect ubiquitin (red) or DAPI (nuclei; blue). Quantification of the percentage of poly-ubiquitin-positive staining in GFP-positive cells (*lower*). Arrowheads indicate the co-localization of the poly-ubiquitin with TDP-43-GFP-positive cells. Data are presented as the means ± SD of 3 independent experiments. **p<0.005, ***p<0.001 (two-way ANOVA with Bonferroni multiple comparison test). Scale bars, 20 μm. **(C)** Immunocytochemistry was subsequently performed to detect the expression of cleaved caspase-3 (red) or DAPI (nuclei; blue). Quantification of the percentage of cleaved caspase-3-positive staining in GFP-positive cells (lower). Arrowheads indicate the co-localization of the caspase-3 with TDP-43-GFP-positive cells. Data are presented as the means ± SD of 3 independent experiments. **p<0.005, ***p<0.001 (two-way ANOVA with Bonferroni multiple comparison test). Scale bars, 200 μm.

### PTK2 inhibition ameliorates UPS impairment and behavioral deficits in *Drosophila* model of TDP-43 proteinopathy

Given strong *in vitro* evidence that PTK2 regulates TDP-43-induced toxicity in neuronal cells, we next examined whether inhibiting PTK2 suppresses TDP-43-induced toxicity *in vivo* using a *Drosophila* model of TDP-43 proteinopathy expressing human TDP-43 and ATXN2-32Q in the nervous system (hereafter referred to as TDP-P fly) (Elden et al, 2010; Kim et al, 2014). The expression of TDP-43 in TDP-P flies was verified by Western blot (Fig. 3A). We first assessed if UPS is impaired in TDP-P flies. To monitor UPS activity in flies, we utilized CL1-GFP reporter flies expressing CL1 fused to GFP. CL1 degron is a peptide that is rapidly degraded specifically by the UPS. Therefore, the level of CL1-GFP is indicative of UPS activity (Pandey et al, 2007). The levels of CL1-GFP protein were markedly increased in TDP-P flies compared to controls (Fig. 3B). Moreover, poly-ubiquitinated proteins were significantly increased in TDP-P fly heads (Fig. 3C), suggesting that UPS is impaired in TDP-P flies.

**Figure 3.**
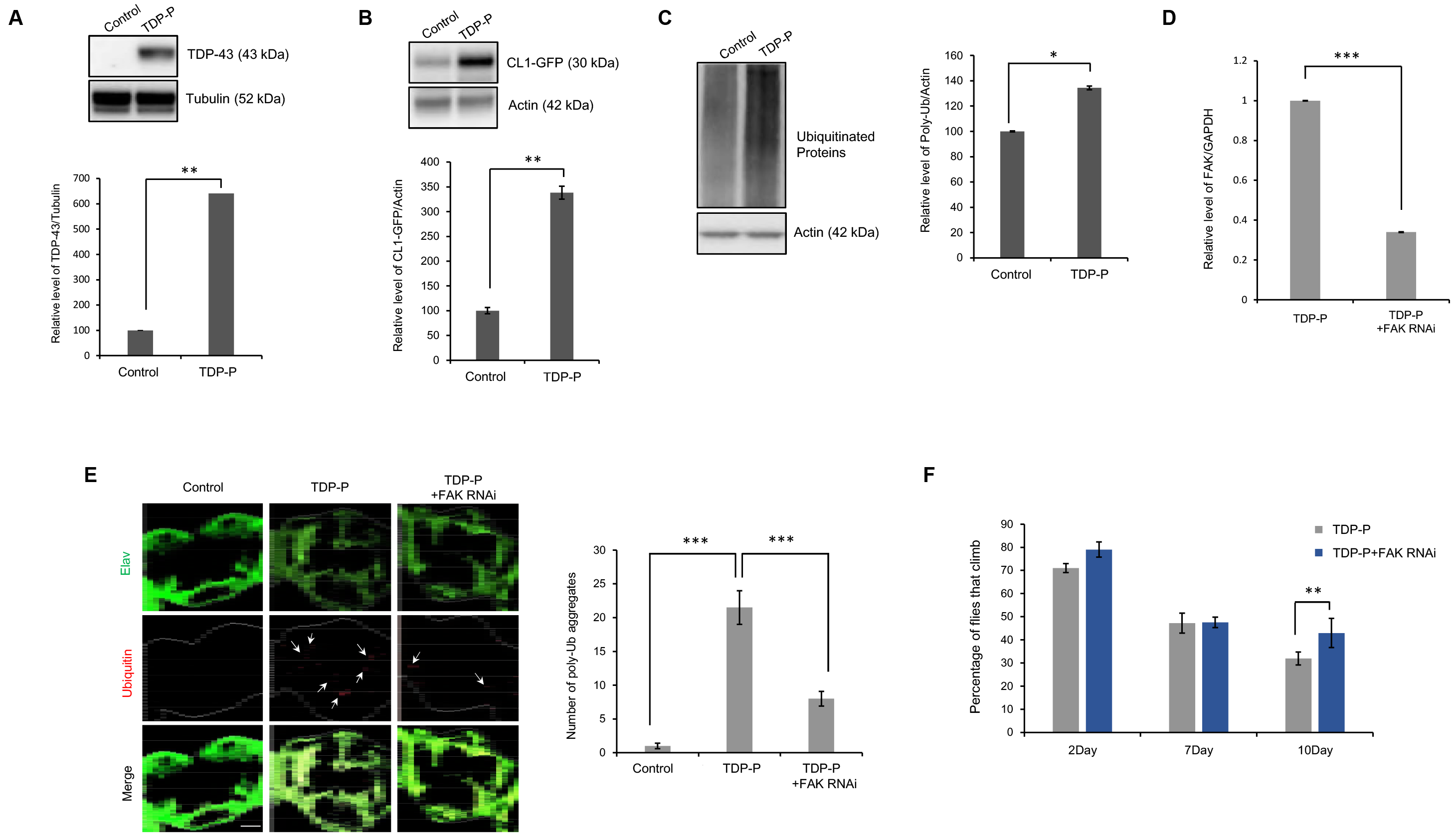
FAK downregulation mitigates UPS impairment and neuronal toxicity of TDP-43-proteinopathy fly model. **(A-C)** Western blot analysis of TDP-43, CL1-GFP, and poly-ubiquitinated proteins from lysates of control or TDP-43-proteinopathy (TDP-P) fly heads. TDP-P expression inhibits UPS activity. Quantification of immunoblots from 3-5 independent experiments and normalized to tubulin or actin. Data are presented as the means ± SEM. *p<0.05, **p<0.005 (Student’s *t*-test). Genotypes: Control is *elavGS*,*UAS-CL1-GFP*/+, TDP-P is *UAS-TDP-43*/+;*elavGS*,*UAS*-*CL1*-*GFP*/*UAS-ATXN2-32Q*. **(D)** RT-PCR for *Fak* mRNA expression from cDNA of TDP-P fly heads. Quantification data of *Fak* mRNA transcript levels are presented as the means ± SD from 3 independent real-time RT-PCR experiments. GAPDH was used for normalization. Genotypes: TDP-P is *UAS-ATXN2-32Q/+;elavGS,UAS-TDP-43/UAS-EGFPRNAi*, TDP-P+FAK RNAi is *UAS-ATXN2-32Q/+;elavGS,UAS-TDP-43/UAS-Fak RNAi*^*MS00010*^. ***p<0.001 (Student’s *t*-test). **(E)** TDP-P or TDP-P+FAK RNAi flies were immunostained for ubiquitin (red) and elav (green). Quantification of the number of poly-ubiquitin-positive inclusions (*right*). Arrowheads indicate poly-ubiquitin-positive inclusions. Data are presented as the means ± SEM of 4 independent experiments. ***p<0.001 (two-way ANOVA with Tukey’s multiple comparison test). Scale bars, 50 μm. Genotypes: Control is *elavGS/+*, TDP-P is *UAS-ATXN2-32Q/+;elavGS,UAS-TDP-43/UAS-EGFP RNAi* and TDP-P+FAK RNAi is *UAS-ATXN2-32Q/+;elavGS,UAS-TDP-43/UAS-Fak RNAf*^*MSm010*^. **(F)** Climbing activity of TDP-P+GFP RNAi and TDP-P+FAK RNAi flies at the indicated timepoints. Quantification of the percentage of flies that climb. Data are presented as the means ± SEM of 4 independent experiments. **p<0.005 (Student’s t-test). Genotypes: TDP-P is *UAS-ATXN2-32Q/+;elavGS,UAS-TDP-43/UAS*-*EGFP RNAi* and TDP-P+FAK RNAi is *UAS-ATXN2-32Q/+;elavGS,UAS-TDP-43/UAS-Fak RNAi*^*HMS000]0*^.

To determine whether inhibiting PTK2 decreases poly-ubiquitinated proteins in TDP-P flies, we inhibited expression of *Fak*, the *Drosophila* homologue of PTK2, by RNAi-mediated gene knock-down (Fig. 3D). Inhibiting FAK dramatically decreased ubiquitin-positive aggregates in TDP-P flies (Fig. 3E). Previously, we revealed that TDP-P flies showed deficits in climbing ability (Elden et al, 2010; Kim et al, 2014). TDP-P flies showed a significant reduction of climbing ability at day 10 (Fig. 3F). When FAK is inhibited, climbing ability was significantly improved in TDP-P flies compared to controls (Fig. 3F). Taken together, PTK2 inhibition ameliorates UPS impairment and behavioral deficits in *Drosophila* model of TDP-43 proteinopathy

### PTK2 inhibition ameliorates TDP-43-induced neurotoxicity in a p62-dependent manner

We next investigated how PTK2 regulates neurotoxicity in TDP-43 proteinopathy. It is noteworthy that inhibiting PTK2 markedly decreased ubiquitinated proteins under the condition that protein degradation via UPS is blocked (Fig. S2B). However, PTK2 inhibition did not restore the impaired proteasome activity in TDP-43 overexpressed neuronal cells (Fig 1D). Theses result suggest that PTK2 inhibition improves the mechanism of ubiquitinated protein degradation other than UPS. Ubiquitinated proteins can be degraded through ALP under the condition of UPS dysfunction (Korolchuk et al, 2010). p62, also known as sequestosome 1 (SQSTM1), is an ubiquitin-binding protein that mediates degradation of poly-ubiquitinated proteins via ALP (Demishtein et al, 2017; Lippai & Low, 2014). To see if p62 is involved in PTK2-mediated neurotoxicity under the condition of UPS dysfunction, we treated PTK2 inhibitor to MG132-treated cells with/without suppressing p62 by p62-specific siRNA (Fig. 4A). Suppressing p62 completely abolished neuroprotective effect of PTK inhibition against MG132-induced toxicity in both N2a cells and primary neurons (Fig. 4B and C).

**Figure 4.**
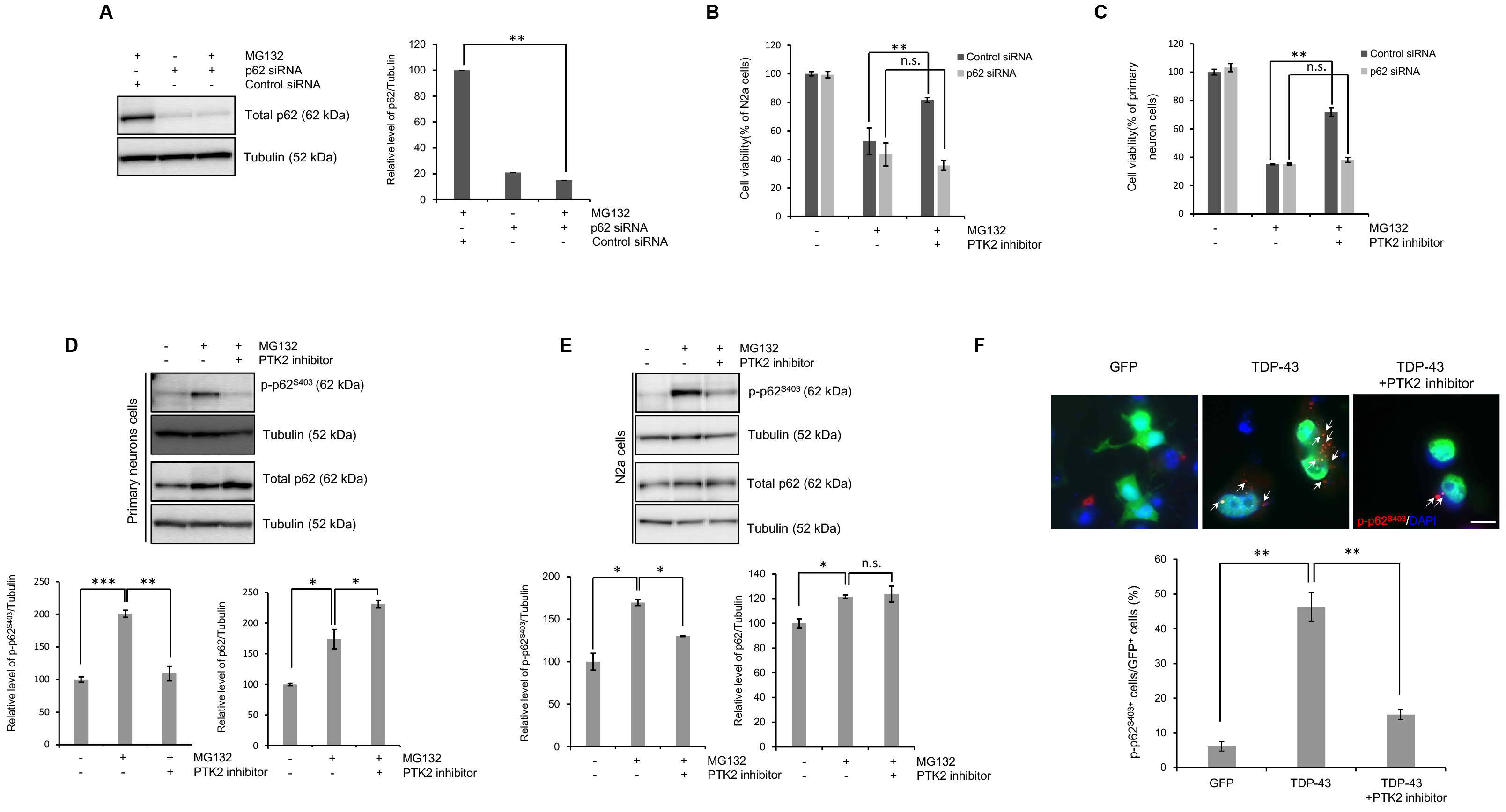
p62 is a crucial component of PTK2-mediated neuroprotection in conditions of UPS impairment. **A-C**, N2a or primary cortical neuron cells were pretransfected with control siRNA (40 nM) or *p62*-specific siRNA (40 nM) for 24 h and subsequently treated with MG132 (5 μm) for 24 h. **(A)** Western blot analysis for total p62 protein expression. Quantification of immunoblots from 3-5 independent experiments and normalized to tubulin. Data are presented as the means ± SD. **p<0.005 (Student’s *t*-test). **(B-C)** A CCK-8 assay was performed to assess the cell viability of p62 inhibition in N2a or primary cortical neuron cells. Data are presented as the means ± SD of 3 independent experiments. **p<0.005, n.s. not significant (one-way ANOVA with Bonferroni multiple comparison test). **D-E**, Primary cortical neuron or N2a cells were pretreated with the PTK2 inhibitor (primary cortical neurons; 0.05 and N2a; 5 μm) for 30 min, subsequently treated with MG132 (5 μm) for 24 h. Western blot analysis for phosphorylated p62 at Ser403 (p-p62^S403^) or total p62 protein expression. Quantification of the immunoblots from 3-5 independent experiments and normalized to tubulin. Data are presented as the means ± SD. *p<0.05, **p<0.005, ***p<0.001, n.s. not significant (one-way ANOVA with Bonferroni multiple comparison test). **(F)** N2a cells were transiently transfected with the control vector (pCMV6-AC-GFP) or TDP-43-expressing construct (pCMV6-AC-TDP-43-GFP) for 2 days and subsequently treated with PTK2 inhibitor (5 μm) for 24 h. Immunostaining was performed thereafter. Immunocytochemistry was subsequently performed to detect p-p62^S403^ (red) or DAPI (nuclei; blue). Arrowheads indicate p-p62^S403^-positive inclusions. Quantification of the percentage of p-p62^S403^-positive staining in GFP-positive cells (lower). Data are presented as the means ± SD of 3 independent experiments. **p<0.005 (one-way ANOVA with Bonferroni multiple comparison test). Scale bars, 10 μm.

We next investigated whether PTK2 regulates p62 under the condition of UPS dysfunction. Inhibiting PTK2 dramatically decreased the levels of phosphorylated p62 at Ser403 (p-p62^S403^) without altering total p62 levels in N2a cells and primary neurons treated with MG132 (Fig. 3D and E). PTK2-mediated regulation of p62 phosphorylation under the condition of UPS dysfunction is conserved in human neuronal cells and specific to Ser403 (Fig. S5A-C). PTK2 did not regulate p62 phosphorylation under basal condition (Fig. S5D and E), suggesting that PTK2-mediated regulation of p62 phosphorylation at Ser403 is specific to the condition of UPS dysfunction. To see if overexpressing TDP-43 also increases p-p62^S403^ levels, we stained N2a cells expressing TDP-43 with p-p62^S403^-specific antibody. Furthermore, PTK2 inhibition effectively reduced the increase of p-p62^S403^ induced by TDP-43 overexpression (Fig. 4F).

The phosphorylation of p62 at Ser403 modulates its binding affinity to poly-ubiquitinated proteins, thereby regulating their autophagic degradation (Matsumoto et al, 2011). To examine the effect of phosphorylation of p62 at Ser403 on the degradation of ubiquitinated proteins under the condition of UPS impairment, we generated stable N2a cell lines expressing wild type p62 and non-phosphorylated form of p62 (p62^S403A^) (Fig. S6A and B). p62^S403A^-expressing cells showed significantly less poly-ubiquitin aggregates than p62-expressing cells under the condition of UPS impairment (Fig. 5A), which was further validated by Western blot analyses (Fig. S6C and D). Moreover, p62^S403A^-expressing cells showed significantly less co-localization of poly-ubiquitin aggregates with autophagic vesicles than p62-expressing cells (Fig. 5A). Consistent with these results, p62^S403A^-expressing cells showed more co-localization of p62 puncta with poly-ubiquitin aggregates than p62-expressing cells (Fig. 5B), indicating that p62 phosphorylation at Ser403 inhibits degradation of poly-ubiquitinated proteins via ALP.

**Figure 5.**
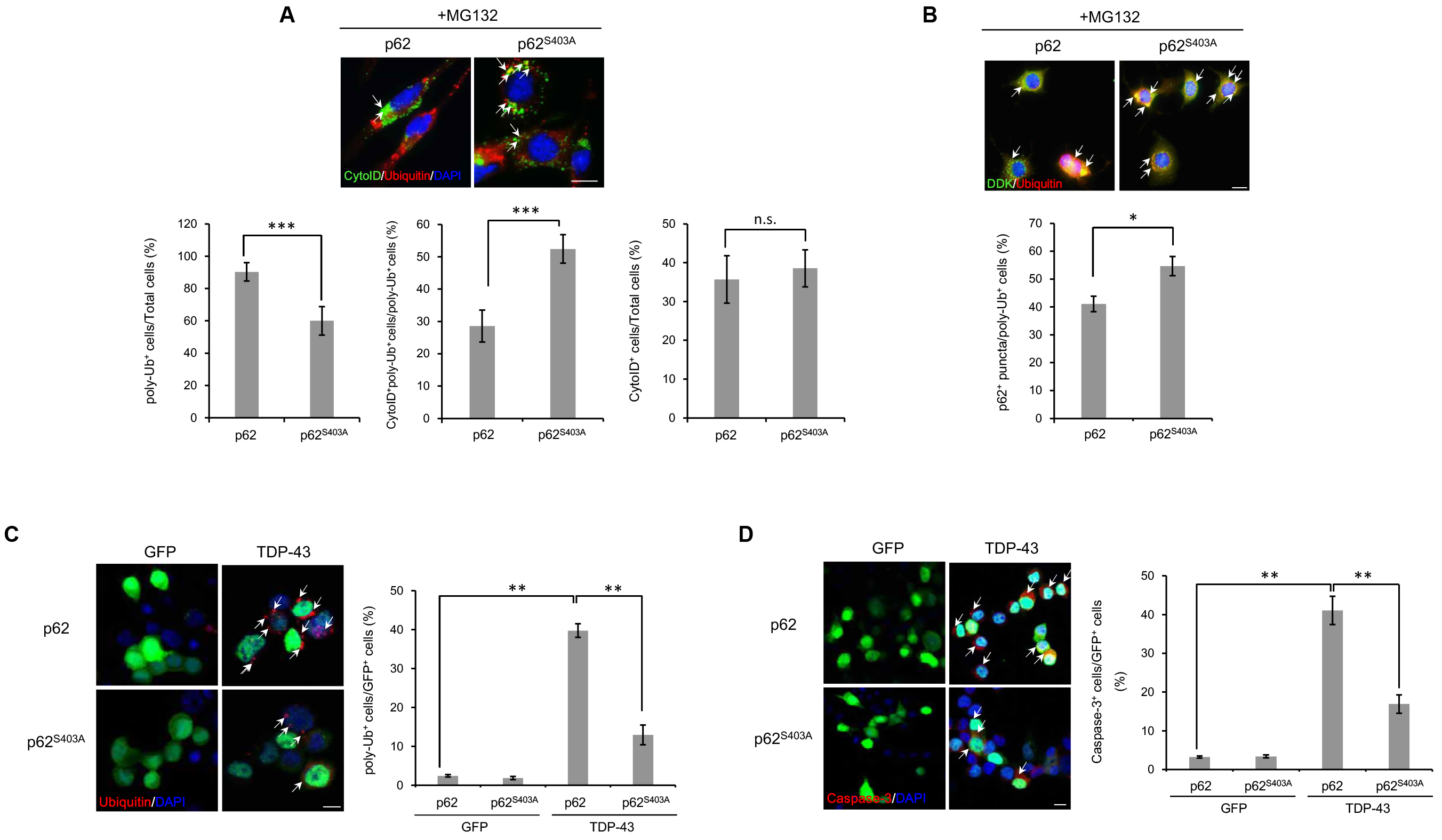
S403A-p62 suppresses TDP-43-and UPS impairment-induced neuronal toxicity in neuronal cells. **A-B**, The p62-, and p62^S403A^-overexpressing N2a stable cells lines treated with MG132 (5 μm) for 24 h. Immunocytochemistry was performed thereafter. **(A)** Immunocytochemistry to detect the Cyto-ID autophagy dye (green) or ubiquitin (red) was subsequently performed. Quantification of the percentage of poly-ubiquitin-positive staining in all cells (*left*), the percentage of CytoID-positive and poly-ubiquitin-positive staining in poly-ubiquitin-positive cells (*middle*), or the percentage of CytoID-positive staining in total cells (*right*). Arrowheads indicate the colocalization of the Cyto-ID autophagy dye with poly-ubiquitin in the p62 stable cells lines. Data are presented as the means ± SD of 3 independent experiments. ***p<0.001, n.s. not significant (Student’s *t*-test). Scale bars, 10 μm. **(B)** Immunocytochemistry to detect the DDK (p62; green) or poly-ubiquitin (red) was subsequently performed. Quantification of the percentage of p62-positive puncta in poly-ubiquitin-positive cells (lower). Arrowheads indicate the co-localization of p62 with poly-ubiquitin in the p62 stable cells lines. Data are presented as the means ± SD of 3 independent experiments. *p<0.05 (Student’s *t*-test). Scale bars, 10 μm. **C-D**, p62-or p62^S403A^-overexpressing N2a stable cells lines were transiently transfected with the control vector (pCMV6-AC-GFP) or TDP-43-expression construct (pCMV6-AC-TDP-43-GFP) for 3 days. Immunostaining was performed thereafter. **(C)** Immunocytochemistry was subsequently performed to detect poly-ubiquitin (red) or DAPI (nuclei; blue). Arrowheads indicate poly-ubiquitin-positive inclusions. Quantification of the percentage of poly-ubiquitin-positive staining in GFP-positive cells (*right*). Data are presented as the means ± SD of 3 independent experiments. **p<0.005 (two-way ANOVA with Bonferroni multiple comparison test). Scale bars, 10 μm. **(D)** Immunocytochemistry to detect caspase-3 (red) or DAPI (nuclei; blue) was subsequently performed. Arrowheads indicate caspase-3-positive cells. Quantification of the percentage of caspase-3-positive staining in GFP-positive cells (right). Data are presented as the means ± SD of 3 independent experiments. **p<0.005 (two-way ANOVA with Bonferroni multiple comparison test). Scale bars, 10 μm.

We next investigated whether p62 phosphorylation at Ser403 regulates TDP-43-induced neurotoxicity. To do this, we transfected TDP-43 or control to N2a cells expressing p62 or p62^S403A^. Accumulation of poly-ubiquitinated aggregates by TDP-43 overexpression was markedly repressed in p62^S403A^-expressing cells compared to p62-expressing cells (Fig. 5C). In line with these, induction of caspase 3 expression by TDP-43 overexpression was strongly suppressed in p62^S403A^-expressing cells compared to p62-expressing cells (Fig. 5D). Taken together, our data suggest that PTK2 inhibition ameliorates TDP-43-induced neurotoxicity in a p62-dependent manner by repressing phosphorylation of p62 at Ser403.

### PTK2 regulates p62 phosphorylation at Ser403 via TBK1

PTK2 is a tyrosine kinase which cannot phosphorylate p62 at Ser403. Therefore, another kinase should be involved in PTK2-mediated regulation of p62 phosphorylation. Previous studies showed that Ser403 of p62 is phosphorylated by TANK-binding kinase 1 (TBK1), casein kinase 2 (CK2), or unc-51 like autophagy-activating kinase 1 (ULK1) (Lim et al, 2015; Matsumoto et al, 2011; Pilli et al, 2012). Among those, only inhibiting TBK1 with BX797 rescued MG132-mediated neurotoxicity in SH-SY5Y cells (Fig. 6A). TBK1 inhibition with TBK1-specific siRNA showed similar protective effect against MG132-mediated neurotoxicity in primary neurons (Fig. 6B and C). TBK1 inhibition repressed MG-132-mediated induction of p62^S403^ levels in primary neurons, without altering the levels of total p62, total PTK2, and p-PTK2^Y397^ (Fig. 6D and E). Although it is not clear how PTK2 regulates TBK1 activity in neurons, out data suggest that PTK2 regulates neurotoxicity induced by UPS impairment by regulating TBK1-p62 axis in neurons.

**Figure 6.**
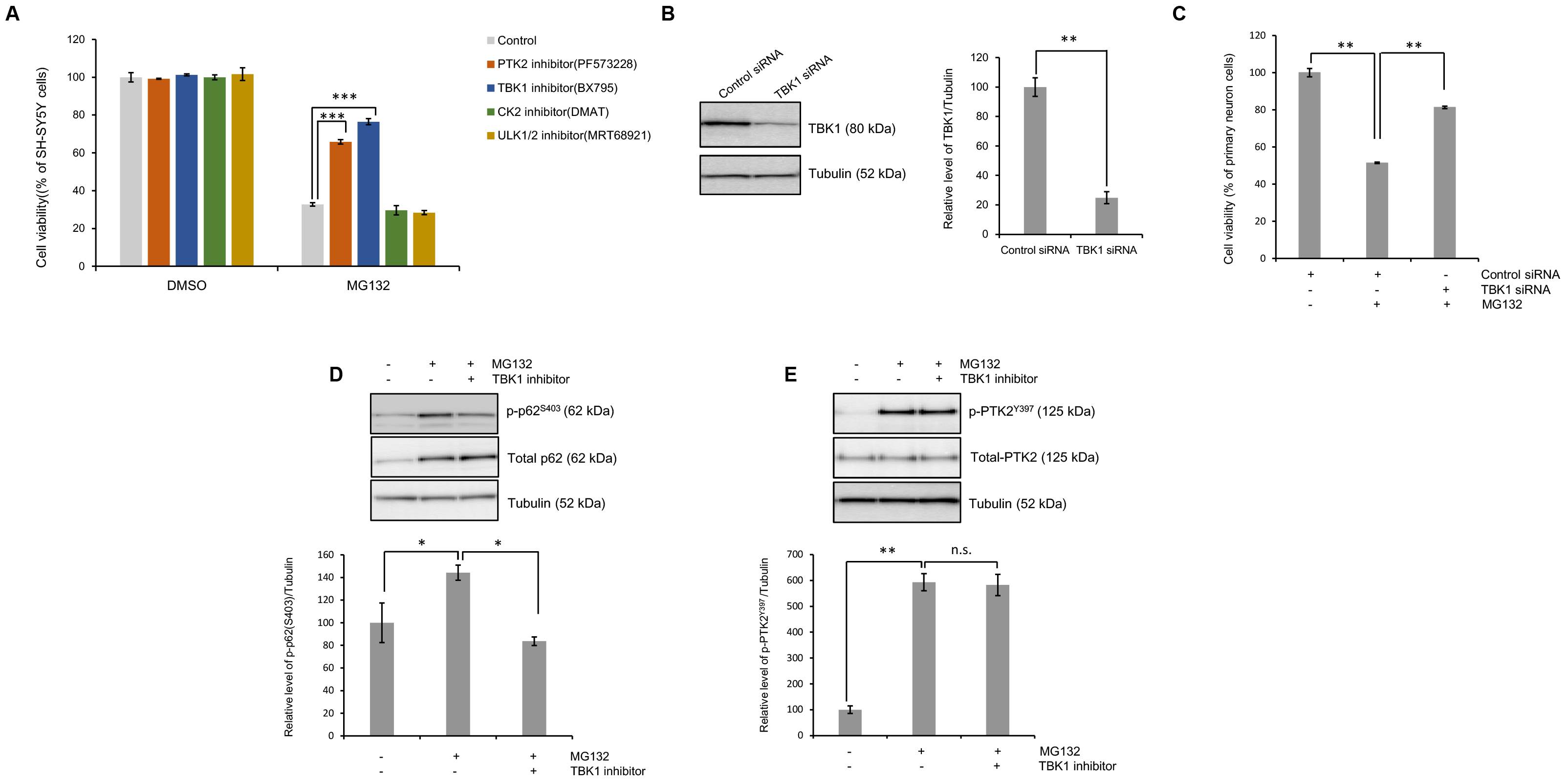
TBK1 inhibition attenuates UPS impairment-induced neuronal toxicity. **(A)** SH-SY5Y cells were pretreated with the PTK2 inhibitor (PF573228; 5 IμM), TBK1 inhibitor (BX795; 1 μm), CK2 inhibitor (DMAT; 5 μm), or ULK1/2 inhibitor (MRT689211; 5 nM) for 30 min and subsequently treated with MG132 (5 μm) for 24 h. CCK-8 analysis was performed thereafter. Data are presented as the means ± SD of 3 independent experiments. ***p<0.001 (one-way ANOVA with Bonferroni multiple comparison test). **B-E**, Primary cortical neuron cells were transfected with the *tbk1*-specific siRNA (100 nM) for 30 min and subsequently treated with MG132 (5 μm) for 24 h. Immunoblotting and CCK-8 analysis was performed thereafter. **(B)** Western blot analysis for TBK1 protein expression. Quantification of the immunoblots from 3-5 independent experiments and normalized to tubulin. Data are presented as the means ± SD. **p<0.005 (Student’s *t*-test). **(C)** A CCK-8 assay was performed to assess the cell viability of TBK1 inhibition in primary cortical neuron cells. Data are presented as the means ± SD of 3 independent experiments. **p<0.005 (one-way ANOVA with Bonferroni multiple comparison test). **(D)** Western blot analysis for p-p62^S403^ or total p62 protein expression. Quantification of the immunoblots from 3-5 independent experiments and normalized to tubulin. Data are presented as the means ± SD. *p<0.05 (one-way ANOVA with Bonferroni multiple comparison test). **(E)** Western blot analysis for p-PTK2^Y397^ or total PTK2 protein expression. Quantification of the immunoblots from 3-5 independent experiments and normalized to tubulin. Data are presented as the means ± SD. **p<0.005, n.s. not significant (one-way ANOVA with Bonferroni multiple comparison test).

Taken together, our data support a model whereby TDP-43 compromises UPS activity (Fig. 7). In the context of TDP-43 proteinopathy, accumulation of TDP-43 could compromise the UPS. UPS impairment eventually leads to increase of ubiquitinated aggregates and PTK2 activation. PTK2 activation also increase TBK1 activity which phosphorylates Serine 403 site of p62. Increased serine 403 phosphorylation of p62 weakens the ability of p62 which transfer ubiquitinated proteins to autophagosome. Consequently, elevated proteotoxic stress by accumulation of ubiquitinated aggregates induces neuronal cell death. On the other hand, PTK2 inhibition reduces the S403 phosphorylation of p62, which enables ubiquitinated proteins to be more effectively degraded through ALP, thereby mitigating TDP-43 toxicity.

**Figure 7.**
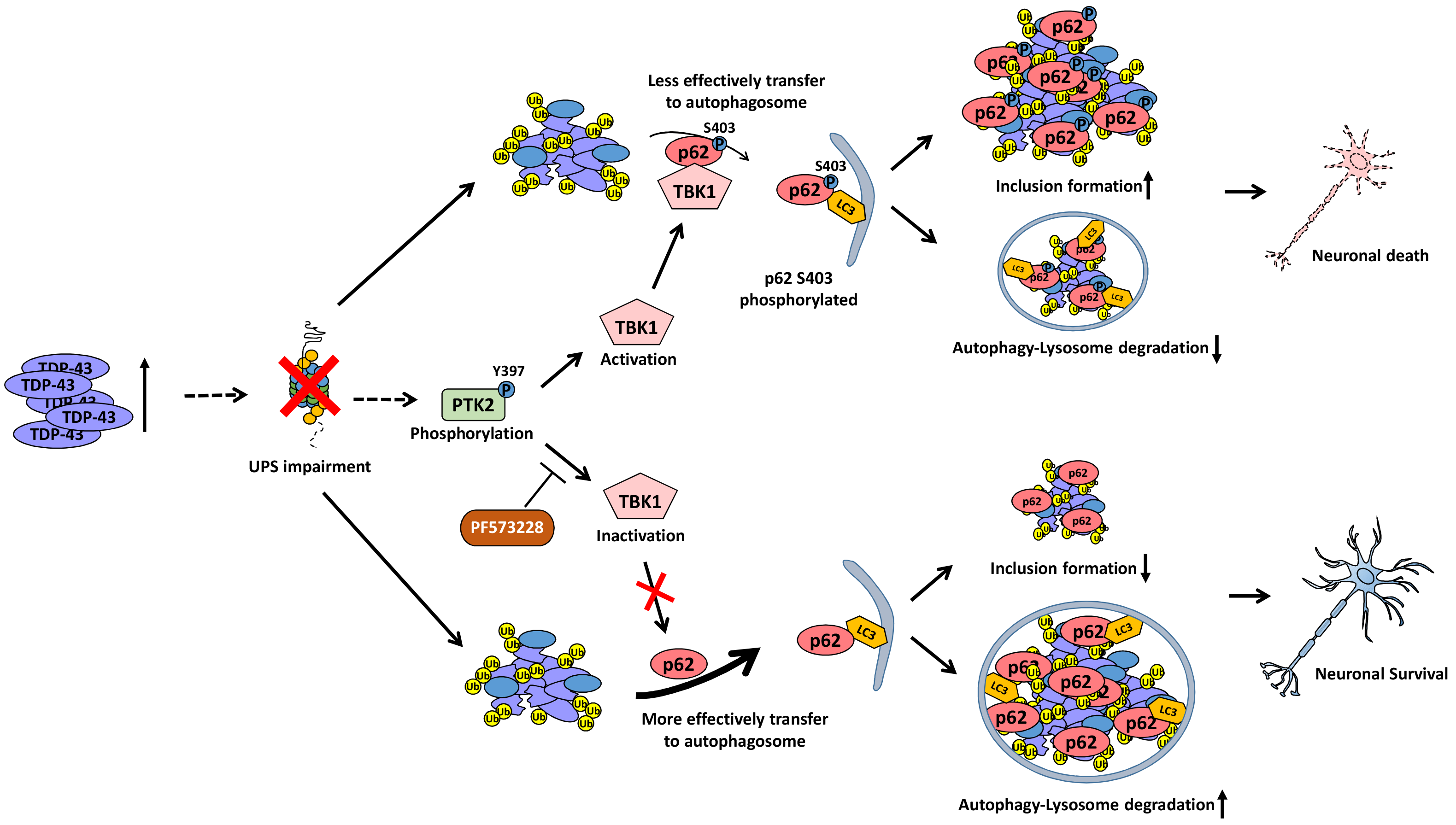
Model of TDP-43 mediated neurodegeneration in TDP-43 proteinopathy. See text for details.

## Discussion

More than 95% of ALS patients have TDP-43 aggregates in their affected tissues (Scotter et al, 2015). As aforementioned, several lines of evidence suggested that UPS may be impaired in ALS with TDP-43 proteinopathy (Chang & Monteiro, 2015; Maekawa et al, 2009; Tashiro et al, 2012; Watanabe et al, 2010). Some evidence suggests that ubiquitin proteasome activity is reduced in the motor neurons of sporadic ALS patients (Bendotti et al, 2012). However, it has never been clearly addressed whether TDP-43 is the culprit behind UPS dysfunction in ALS. In this study, we demonstrated that TDP-43 overexpression impairs UPS in cellular and *Drosophila* models of TDP-43 proteinopathy. In particular, 26S proteasomal activity is significantly reduced by TDP-43 overexpression. Our data suggest that abnormal accumulation of TDP-43 contributes to the progression of ALS with TDP-43 proteinopathy at least in part via impairing UPS. Further studies are warranted to determine how TDP-43 accumulation affects UPS function.

Through kinase inhibitor screening in neuronal cells, we identified PTK2 as a potent modifier of toxicity induced by UPS impairment. We demonstrated that PTK2 inhibition suppresses neurotoxicity induced by UPS impairment. Moreover, PTK inhibition represses the accumulation of ubiquitinated aggregates in cellular and *Drosophila* models of TDP-43 proteinopathy, suggesting that inhibiting PTK2 may represent a novel therapeutic intervention for neurodegenerative diseases with TDP-43 proteinopathy.

p62 mediates sequestering poly-ubiquitinated proteins into the autophagosome (Bjorkoy et al, 2005; Pankiv et al, 2007). Previous studies showed that p62 plays an important role in the degradation of poly-ubiquitinated proteins under the condition of UPS impairment (Demishtein et al, 2017; Kageyama et al, 2014; Milan et al, 2015; Pankiv et al, 2007). In this study, we demonstrated that p-p62^S403^ levels are increased by UPS impairment and TDP-43 overexpression in neuronal cells. PTK2 inhibition represses induction of p-p62^S403^ upon UPS impairment in neuronal cells. Expressing non-phosphorylated form of p62 (p62^S403A^) increased the co-localization of ubiquitin aggregates with autophagic vesicles and decreased accumulation of ubiquitinated aggregates induced by TDP-43, indicating that p62^S403A^ facilitates the degradation of ubiquitinated proteins via ALP in UPS impaired condition in neuronal cells. Importantly, expressing non-phosphorylated form of p62 (p62^S403A^) significantly attenuated cell death induced by TDP-43 overexpression in neuronal cells. Taken together, our findings suggest that PTK2 regulates TDP-43-mediated neurotoxicity by modulating phosphorylation of p62 at Ser403. Intriguingly, mutations in *SQSTM1* are associated with ALS and FTLD (Le Ber et al, 2013; Rubino et al, 2012; Shimizu et al, 2013; Teyssou et al, 2013; Yang et al, 2015). In addition, p-p62^S403^ positive inclusions are accumulated in the spinal cord of ALS patients (Kurosawa et al, 2016). These results raise the possibility that PTK2-p62 axis plays a critical role in the pathogenesis of ALS.

TBK1 is a kinase which phosphorylates p62 at Ser403 (Matsumoto et al, 2015). Of note, mutations in TBK1 are also associated with ALS (Cui et al, 2018). Moreover, optineurin (OPTN), another interacting partner of both TBK1 and p62, is associated with ALS (Markovinovic et al, 2017). TBK1 phosphorylates OPTN to modulate its binding affinity with ubiquitin chain (Richter et al, 2016). ALS-associated mutations in *OPTN* are reported to induce deficits in autophagosome formation and thereby inhibit degradation of ubiquitinated substrates (Bansal et al, 2018; Chang & Monteiro, 2015; Sundaramoorthy et al, 2017). Taken together, all the evidence suggests that UPS dysfunction may be one of core pathogenic mechanisms of ALS. We demonstrated that TBK1 inhibition strongly suppresses induction of p-p62^S403^ and cell death induced by UPS impairment in neuronal cells, suggesting that PTK2-TBK1-p62 axis may represent a novel therapeutic target for ALS. Further studies warranted to elucidate the underlying mechanism how TDP-43 accumulation affects PTK2-TBK1-p62 axis.

In summary, we demonstrated that UPS is impaired in TDP-43 proteinopathy. We identified that PTK2-TBK1-p62 axis regulates degradation of ubiquitinated proteins and neurotoxocity induced by TDP-43 overexpression. Therefore, our data suggest that targeting PTK2-TBK1-p62 axis may represent a novel therapeutic intervention for neurodegenerative diseases with TDP-43 proteinopathy.

## Author Contributions

S.L., Y.-M.J., S.K., Y.K., M.J., and Y.-N.J., planned and performed the experiments.
S.L., S.R.K., K.J.L., J.K., and S.B.L. provided ideas for the project and participated in writing the paper. S.L., K.K., and H.-J.K. wrote the paper.

## Acknowledgements

This work was supported by the KBRI Research Program of the Ministry of Science, ICT & Future Planning (18-BR-02-06, 18-BR-02-03, and 18-BR-01-01); the DGIST R&D Program of the Ministry of Science, ICT and Future Planning (16-BD-0402) (S.B.L.); the Soonchunhyang University Research Fund; the KBSI research fund (D37627 to S.L.); the Basic Science Research Program through the National Research Foundation of Korea (NRF), funded by the Ministry of Science, ICT & Future Planning (NRF-2017R1C1B2007941, NRF-2017R1C1B1008825, NRF-2015R1D1A1A01059079); and the Korea Health Technology R&D Project through the Korea Health Industry Development Institute (KHIDI), funded by the Ministry of Health and Welfare, South Korea (grant number: H I14C1135, HI15C1928). Confocal microscopy data were acquired in the Advanced Neural Imaging Center in KBRI.

## Abbreviations

ALS: Amyotrophic lateral sclerosis
FTLD: Frontotemporal lobar degeneration
UPS: Ubiquitin proteasome system
ALP: Autophagy lysosomal pathway
SQSTM1: Sequestosome 1
TBK1: TANK-binding kinase 1
CK2: Casein kinase 2
ULK1: Unc-51 like autophagy-activating kinase 1
SOD1: Superoxide dismutase 1
OPTN: Optineurin

